# Effects of using deep learning to predict the geographic origin of barley genebank accessions on genome-environment association studies

**DOI:** 10.1101/2025.02.13.638161

**Authors:** Che-Wei Chang, Karl Schmid

**Affiliations:** University of Hohenheim, Stuttgart, Germany

**Keywords:** local adaptation, genome-environment association, barley

## Abstract

Genome-environment association (GEA) is an approach for identifying adaptive loci by combining genetic variation with environmental parameters, offering potential for improving crop resilience. However, its application to genebank accessions is limited by missing geographic origin data. To address this limitation, we explored the use of neural networks to predict the geographic origins of barley accessions and integrate imputed environmental data into GEA. Neural networks demonstrated high accuracy in cross-validation but occasionally produced ecologically implausible predictions as models solely considered geographical proximity. For example, some predicted origins were located within non-arable regions, such as the Mediterranean Sea. Using barley flowering time genes as benchmarks, GEA integrating imputed environmental data (N=11,032) displayed partially concordant yet complementary detection of genomic regions near flowering time genes compared to regular GEA (N=1,626), highlighting the potential of GEA with imputed data to complement regular GEA in uncovering novel adaptive loci. Also, contrary to our initial hypothesis anticipating a significant improvement in GEA performance by increasing sample size, our simulations yield unexpected insights. Our study suggests potential limitations in the sensitivity of GEA approaches to the considerable expansion in sample size achieved through predicting missing geographical data. Overall, our study provides insights into leveraging incomplete geographical origin data by integrating deep learning with GEA. Our findings indicate the need for further development of GEA approaches to optimize the use of imputed environmental data, such as incorporating regional GEA patterns instead of solely focusing on global associations between allele frequencies and environmental gradients across large-scale landscapes.

## Introduction

Crop domestication and improvement have led to genetic bottlenecks, resulting in reduced genetic diversity in modern crop cultivars compared to their wild progenitors (Khoury et al. 2022; Tanksley and McCouch 1997). The erosion of genetic diversity poses a significant challenge for contemporary crop breeding, particularly in view of climate change that threatens agricultural productivity and resilience. To address the loss of adaptive genetic variation during domestication, plant breeders seek novel genetic variation in exotic genetic resources, such as traditional landraces and wild relatives (Bohra et al. 2022; Kumar et al. 2020). However, the introduction of useful alleles from these resources is hindered by linkage drag and potential reproductive barriers (Bohra et al. 2022; Dempewolf et al. 2017; Saad et al. 2022). In the genomics era, the integration of advanced sequencing technologies and computational approaches provides a promising avenue for introgressing favorable traits by mapping and selection of beneficial alleles using molecular breeding methods.

Genome-environment association (GEA), or environmental genome-wide association (EnvG-WAS) studies, identify candidate adaptive loci associated with local adaptation by correlating allele frequencies with environmental variables (Coop et al. 2010; Lasky et al. 2012, 2015). GEA studies are based on the assumption that plants adapt to their local environments through natural selection, which drives changes in allele frequencies at causal genes influencing environmental adaptation across different regions in the distribution range of a species (Lasky et al. 2023). The application of GEA to crops and their wild relatives, such as sorghum (Lasky et al. 2015), rice (Gutaker et al. 2020), wild tomato (Gibson and Moyle 2020), sunflower (Todesco et al. 2022), and barley (Russell et al. 2016), indicates its potential to facilitate the identification and introgression of useful genetic variation into advanced breeding populations.

GEA approaches require a large number of geo-referenced samples from diverse environments to extract environmental data, whereas collecting such samples from scratch is costly (Campbell et al. 2025). In contrast, genebanks preserve genetic diversity through extensive exsitu collections of traditional crop cultivars, and they serve important sources of novel and useful genetic variation for inclusion into plant breeding programs (Mascher et al. 2019). However, a significant challenge lies in efficiently identifying valuable genetic variants among thousands of genebank accessions, particularly given the limited resources available for their evaluation (Longin and Reif 2014; Schulthess et al. 2022). Recent advancements in affordable high-throughput sequencing have allowed to unlock novel genetic diversity (Mascher et al. 2019) and have enabled comprehensive genomic analyses of genebank collections for various crops, including barley (Milner et al. 2019), wheat (Sansaloni et al. 2020; Schulthess et al. 2022), rice (Gutaker et al. 2020; Wang et al. 2018), pepper (Tripodi et al. 2021), and chickpea (Varshney et al. 2021).

Despite the rapid increase in genome-wide sequencing data, a relatively small proportion of genebank collections has a detailed record of their geographical origin, as many were collected before the establishment of modern collection data standards. For instance, among the 20,000 barley accessions from the German genebank at IPK that were genotyped by sequencing (Milner et al. 2019), only about 13% are geo-referenced (König et al. 2020). This limitation hinders the application of computational genome-environment association (GEA) approaches to identify adaptive genetic variants in plant genetic resources. However, the advent of deep learning methods, such as neural networks, along with advancements in computer hardware like graphics processing units (GPUs), has enabled computationally efficient inference of geographical origins for large datasets. For example, Battey et al. (2020) developed *Locator*, a neural network-based tool that predicts geographical origins using complex allele distribution patterns. It outperforms conventional modelbased approaches, such as SPASIBA (Guillot et al. 2016), in both computational efficiency and prediction accuracy. These advancements create the opportunity to recover missing geographical origin information for genebank accessions directly from genome sequencing data. The inferred geographic coordinates for additional accessions can subsequently be used in GEA, potentially enhancing the statistical power of environmental association studies by leveraging a much larger sample size (Cockram and Mackay 2018).

In this study, we explore the potential of applying geographical origin inference using the Locator method by Battey et al. (2020) to improve the detection of adaptive loci reflecting selection by environmental factors using GEA approaches in genebank collections. We refer to this approach as *GEAplus* framework and apply it using the data of the global barley collection (Milner et al. 2019). Barley is one of the most important crops worldwide and its data set is suitable for this approach because of its large size with more than 10,000 genotyped accessions. We first assess the prediction accuracy of geographical origin inference by deep learning in barley landraces and compare the results of conventional GEA with those of *GEAplus* to test the hypothesis that a larger sample leads to the discovery of additional genomic regions involved in local adaptation. To validate the observed results *GEAplus*, we perform conventional GEA and *GEAplus* on populations generated through forward-in-time simulations of genetic variation under different models of domestication, and then compare the power of conventional GEA and *GEAplus* to identify causal single nucleotide polymorphisms (SNPs) affecting fitness in different environments.

## Materials and Methods

### Overview of *GEAplus* framework

We developed the *GEAplus* framework (Figure 1) to investigate genomic adaptation by using genebank samples that lack geographical origin data. The framework begins by constructing a fully connected neural network model trained on geo-referenced samples with both geographical origin and genotypic data. This predictive model infers the geographical origins of a prediction set—samples without passport data—using genome-wide genetic variants, without relying on an explicit isolation-by-distance model (Battey et al. 2020). This step involves imputing the geographical origin of accessions based on the spatial distribution of SNPs. Once the geographical origins are inferred, environmental data for the prediction set samples are retrieved from public databases. Finally, genome-wide association studies using a mixed linear model are conducted across all samples, including both the training set and the prediction set, to identify potential adaptive loci by testing associations between genetic variants and environmental variables. For clarity, we referred to the traditional GEA approach using only geo-referenced accessions as *regular GEA* in this work to distinguish it from *GEAplus*, which incorporates predicted geographical coordinates.

**Figure 1.**
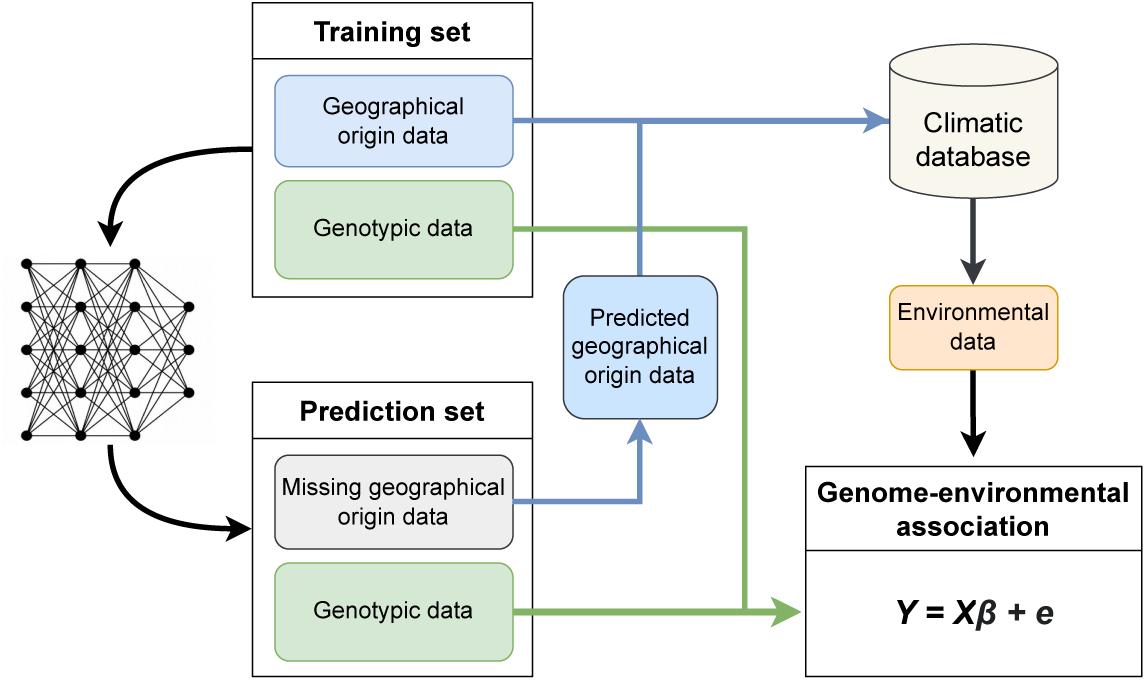
Flowchart of the *GEAplus* framework. Accessions with both geographical origin and genotypic data are used as a training set to build a prediction model using a fully connected neural network (Battey et al. 2020). Once the model is trained, it predicts the geographical origins of accessions lacking passport data based on their genotypic information. Subsequently, environmental data corresponding to the predicted geographical origins are retrieved from public databases. Finally, genome-environment association (GEA) analyses are performed, incorporating genotypic data and various environmental parameters, including those derived through the imputed geographical origins.

### Analysis of a global barley landrace collection

#### Available data of the barley landrace accessions

To evaluate the effectiveness of *GEAplus*, we applied both the regular GEA and *GEAplus* to a global collection of barley landraces. We identified 12,129 landrace accessions from the BRIDGE database (König et al. 2020) by selecting those classified as *Traditional cultivar/landrace*. For the GEA analysis, we retrieved geographical coordinates for 1,661 geo-referenced accessions from the same database (König et al. 2020). As a result, only about 14% of the landrace accessions included in this study had precise geographical origin data. It is important to note that caution should be exercised with the collection site data due to incomplete documentation of seed exchanges or potential errors in passport data, as highlighted in a previous study (Milner et al. 2019).

All selected landrace accessions were genotyped using genotyping-by-sequencing (GBS) by Milner et al. (2019). Single-nucleotide polymorphisms (SNPs) were identified as Milner et al. (2019) with aligning sequences against the ‘Morex’ V3 genome assembly (Mascher et al. 2021). Subsequently, SNPs were filtered using VCFtools v0.1.17 (Danecek et al. 2011) with a minor allele count ≥ 10 and a maximum missing rate < 95%. Missing values were then imputed for all landrace accessions using BEAGLE v5.2 (Browning et al. 2018) with default settings.

In this study, we used the deep learning model *Locator* (Battey et al. 2020) to infer missing geographical origin data. This model predicts geographical origins from genotypic data using a fully connected neural network with a loss function designed to minimize the Euclidean distance between the predicted and true locations of samples in the training data. Simulations in Battey et al. (2020) indicate that *Locator* requires a high density of SNPs to achieve accurate predictions of geographical origin. To meet this requirement, we applied a relaxed filtering criterion, resulting in 557,991 SNPs with a minor allele frequency (MAF) > 0.01 among geo-referenced accessions across the entire landrace collection. For genome-environment association (GEA) analyses, we further filtered these to select 87,036 SNPs with a minor homozygous genotype frequency > 0.01 and a heterozygosity < 0.1 among geo-referenced accessions. Two datasets were prepared with the selected 87,036 SNPs: one for the geo-referenced accessions and another for the entire landrace collection. These datasets were used to perform regular GEA and *GEAplus*, respectively.

We then retrieved bioclimatic data for individual landrace accessions from the WorldClim 2.1 database (Fick and Hijmans 2017). We extracted 19 environmental variables based on the geographical coordinates of origin using the *raster* package (Hijmans 2018). Subsequently, we performed a principal component analysis (PCA) on the 19 environmental variables and used the first three principal components in the downstream GEA analysis to address the multicollinearity of the environmental variables (Hoban et al. 2016).

#### Data cleaning

Due to seed exchange and other historical human activities, the genetic similarity of barley germplasm may conflict with their documented geographical origins (Milner et al. 2019). This disruption of the geo-genetic association weakens the isolation-by-distance pattern, which is expected to reduce the accuracy of geographical origin inference. To enhance prediction accuracy, we removed outliers exhibiting unusual geo-genetic patterns using the *GGoutlieR* R package (version 1.0.2) (Chang and Schmid 2023). As input for *GGoutlieR*, we first calculated ancestry coefficients for the 1,661 geo-referenced accessions using 29,219 representative SNPs. These SNPs were pruned with *PLINK* (Chang et al. 2015) using an *r*^2^ *<* 0.1 threshold. The optimal number of ancestral populations (*K*) was determined with the *estimate_d* function, and ancestry coefficients were estimated using the *alstructure* function from the R package *ALStructure* (Cabreros and Storey 2019). Outlier samples that violated the isolation-by-distance assumption were identified using *GGoutlieR* with the parameter *p_thres = 0.025*, while all other settings were kept at their default values. These outliers were subsequently excluded from the training of the *Locator* model.

#### Inference of geographical origin

We used all accessions that passed the *GGoutlieR* filter to train the neural network model, *Locator* v1.2 (Battey et al. 2020). In assessing the usefulness of geographical origin inference with *Locator*, our main focus was on the general accuracy and robustness, rather than obtaining the most accurate model by tuning the hyperparameters of the neural network model, so the default setting was used.

We evaluated the prediction accuracy using ten-fold cross-validation with ten replicates. The prediction accuracy was measured as the *R*^2^ between a true coordinate and a predicted coordinate. To extract environmental data for GEA analysis (Figure 1), we averaged the predicted coordinates from all prediction models for each accession. To train the model and perform the prediction of geographical origin inference, we used four Nvidia A100 GPUs and 256 Gb working memory of the high-performance computing clusters of the State of Baden-Württemberg (bwHPC; https://www.bwhpc.de/), Germany.

#### GEA analysis with barley landraces

Due to the large sample size, we employed the computationally efficient genome-wide association approach *REGENIE* v2.2.4 (Mbatchou et al. 2021) for GEA analysis. To fit the kinship effect, *REGENIE* uses representative SNPs to conduct genome-wide ridge regression in the first stage of its algorithm. We used a set of SNPs with *r*^2^ *<* 0.2, extracted using *PLINK*, for the genome-wide ridge regression of *REGENIE*. We also used the first three principal components of genotypic data, computed with *PLINK*, for controlling the population structure in the first and second stages of *REGENIE*. To control for false positive associations, we computed adjusted pvalues based on chi-square statistics from *REGENIE* using the genomic inflation factor method (François et al. 2016). Chi-square statistics were divided by an inflation factor *λ*, calculated as *λ* = Median 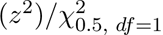, where *z*^2^ is the estimated chi-square statistics and 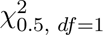 is the middle quantile of a chi-square distribution with one degree of freedom. Adjusted p-values were calculated using the R code of the genomic inflation factor method from Capblancq et al. (2018). We conducted regular GEA and *GEAplus* analysis separately using the geo-referenced accessions and the entire landrace collection. The environmental variables for the entire landrace collection included data from both known geographical origins of geo-referenced samples and predicted origins of non-geo-referenced samples. Since flowering time genes are crucial for barley adaptation (Russell et al. 2016), we used them as benchmarks for successful identification of adaptive loci. To obtain the positions of flowering time genes on the ‘Morex’ V3 genome assembly, we queried the NCBI database (www.ncbi.nlm.nih.gov/) with the following keywords: *heading[All Fields] OR CEN-like[All Fields] OR flowering[All Fields] OR vernalization[All Fields] OR PPD[All Fields] OR VRN-H1[All Fields] AND “Hordeum vulgare”[porgn] AND alive[prop]*. For our analysis, we parsed the physical positions of eleven flowering time genes on the ‘Morex’ V3 genome assembly, including *FT3*, *ELF3*,*PPD-H1*, *FT4*, *CEN*, *FT2*, *FT5*,*TFL1*, *ABAP1*, *VRN-H1*, and *FT1* (reviewed by Fernández-Calleja et al. 2021).

### SLiM simulation

To assess the performance of *GEAplus* framework, we simulated populations under environmental selection with gene flow using *SLiM* v4.0 (Haller and Messer 2023). SLiM is a tool for forward genetic simulations of evolutionary processes with defined parameters, such as mutation rates and migration rates. We implemented a spatial stepping-stone model to approximate gene flow on a continuous landscape, with sub-populations located at real collection sites of barley landraces in the Fertile Crescent and surrounding regions. The simulation focused on a relatively small geographical range instead of a global level to ensure it was computationally feasible. We obtained the geographical coordinates of collection sites from the *Genesys* database (https://www.genesys-pgr.org/) and selected 312 sites with spatial sampling to ensure that each site was at least 40 km away from the others. Since we did not simulate long-distance migration between sites and mislabeling of samples, additional data cleaning with *GGoutlier* was not needed.

We performed simulations under two demographic scenarios. The first scenario involved range expansion from a single refugium (1R) near an archaeological site of domesticated barley in the Israel-Jordan area (Mascher et al. 2016; Sallam et al. 2024) (Figure S1a). The second scenario involved expansion from two refugia (2R), with the second refugium located around the Zagros Mountains based on the hypothesis of a second domestication center (Morrell and Clegg 2007; Figure S1b). The migration rates between adjacent sites per generation were defined as a function of inverse geographical distances. We conducted three replicates of simulations for each demographic scenario by setting different numeric seeds for each replicate. Our simulations were performed with the *nonWF* model of *SLiM*, which supports dynamic population sizes, allowing simulations of sub-populations to propagate from a single individual and enabling the possibility of population extinction.

We ran simulations for 20,000 generations, with each run initiated with a burn-in phase to establish standing variation followed by 10,000 post-burn-in generations as barley was domesticated ∼10,000 years ago (Badr et al. 2000; Sallam et al. 2024). During the burn-in phase, no migration events were allowed, and the carrying capacity of the initial site was set to 5,000 individuals. Population expansion was initiated in the post-burn-in phase, with migration allowed for 10,000 generations. We controlled the total individual number by setting a carrying capacity of 250 individuals for each site in the post-burn-in phase, resulting in approximately 78,000 individuals in our simulation.

We simulated 10^6^ loci for each individual, including 999,900 neutral loci and 100 biallelic selected SNPs, or quantitative trait loci (QTLs), evenly distributed across 10 linkage groups. To compensate for the small individual number, we used the method of Matz et al. (2020) by setting the mutation rate to 6.5 × 10*^−^*7, which is 100-fold higher than the estimation in wild barley (Li et al. 2020). The outcrossing rate was set to 0.01, as the estimated outcrossing rate ranged from 0-1.8% (Abdel-Ghani et al. 2004). The mutation effects of QTLs were drawn from a normal distribution *N* (0, 0.45) (see Supplementary Materials for details). We calculated the phenotype of an individual as the sum of QTL effects and a random value drawn from the normal distribution *N* (0, 0.5) to simulate environmental noise.

To simulate local adaptation, we transformed phenotypes into fitness values for each individual based on local environmental conditions, following the approach described by Haller and Messer (2016). We treated the mean temperature of the warmest quarter from the WorldClim2 (Fick and Hijmans 2017) as an ideal local optimum phenotype for each site. We calculated the relative fitness of an individual as the ratio of its phenotype to the local optimum phenotype. Fitness was defined as the probability density of the normal distribution *N* (*y_opt,j_, σ_plasticity_*), where *y_opt,j_* is the local optimum phenotype for the *j*th site and *σ_plasticity_*is a parameter that controls the strength of selection, which was set to 2.85 (see Supplementary Materials for details). With this setting, the relative fitness of an individual approached to 1 at a given site if its phenotype was close to the locally optimal phenotype.

To reduce the computational demands of our simulations, we adopted a hybrid strategy that combined forward and backward simulations, as described by Haller et al. (2019). In brief, we conducted forward simulations using *SLiM* with the tree-sequence recording method (Haller et al. 2019; Kelleher et al. 2018) to record pedigrees. Neutral mutations were suppressed in the forward simulation and overlaid in the backward simulation according to the tree sequences. This approach ignored neutral mutations in the forward simulation, as neutral mutations were assumed not to affect genealogies (Haller et al. 2019). We then used *pyslim* v1.0.1 and *tskit* v0.5.3 (Kelleher et al. 2018) within a *Python* 3.8 environment to read and manipulate the tree sequences from *SLiM*. We performed the backward simulation of neutral mutations using *msprime* v1.2.0 (Baumdicker et al. 2022) with a mutation rate of 6.5 × 10*^−^*7. At the end of our simulations, we sampled 50 individuals from each site and recorded their genotypes, resulting in a VCF file with 15,600 individuals. We removed loci with MAF less than 0.005 using *vcftools* v0.1.17 (Danecek et al. 2011) as we were not interested in rare alleles. The scripts of simulations are available at https://gitlab.com/kjschmidlab/geaplus.

### Accuracy of geographical origin inference

To simulate the absence of geographical origin data for genebank accessions, we masked the geographical origin information for 95% of individuals in the simulated dataset. First, we randomly selected 50% of the simulated sub-populations (156 out of 312) for further sampling, reflecting the fact that genebank accessions may originate from populations without recorded geographical coordinates, such as those collected before the advent of the Global Positioning System (GPS). Next, we assumed that geographical origin records were largely missing for accessions from the selected sub-populations. Based on this assumption, we randomly selected five individuals from each chosen site to form a training set and masked the geographical origins of the remaining individuals to create a prediction set. By repeating this sampling procedure, we generated three replicates of training sets (*N* = 780) and prediction sets (*N* = 14, 820) for each simulation run.

To infer the missing geographical origins, we employed the *Locator* deep neural network model (Battey et al. 2020). To prevent bias in the predictions, QTLs were excluded from the *Locator* model during training, ensuring they were not used to predict geographical coordinates. For model training, the individuals in the training set were randomly divided into ten groups, with one group sequentially designated as the validation set in each iteration. This procedure was repeated eight times, resulting in a total of 80 trained models for each training set. Following prediction, the geographical origins inferred by the 80 models for each masked individual were averaged to produce the final estimates. The accuracy of geographical origin inference was assessed using the *R*^2^ value, comparing the true locations with the predicted locations.

Since 50% of the simulated sub-populations were excluded from the model training, individuals originating from these removed sub-populations were labeled as Type 1 samples (Figure 2) for clarity. Conversely, individuals sampled from sub-populations included in the model training were categorized as Type 2 samples (Figure 2). Predicting origins for Type 2 samples reflects a scenario where samples with unknown origins might actually belong to a population present in the training set, introducing the potential for data leakage in prediction models (Bernett et al. 2024). To evaluate performance, we computed the *R*^2^ separately for Type 1 and Type 2 samples, alongside measuring prediction errors as the geographical distance between true and predicted locations.

**Figure 2.**
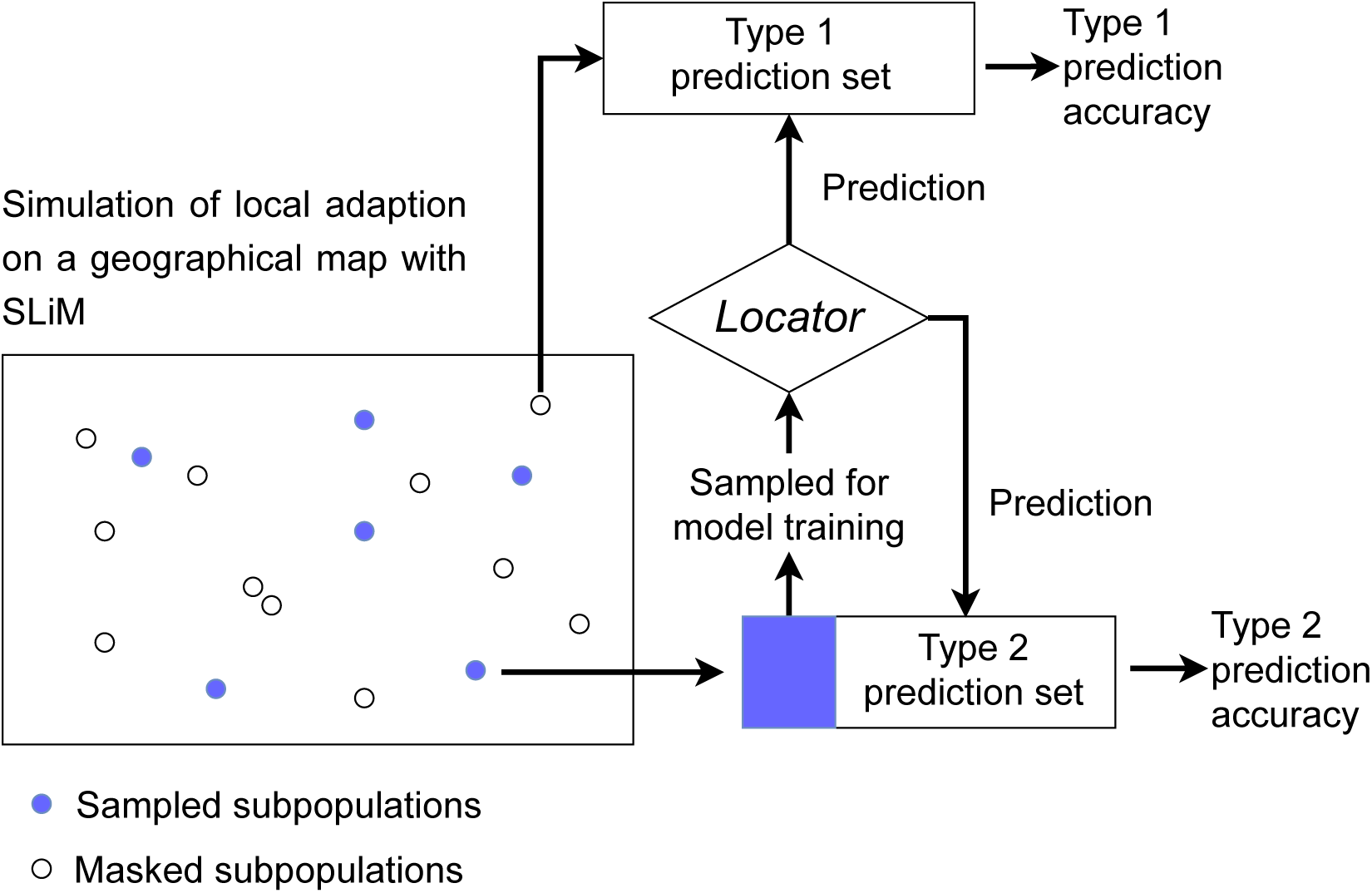
Design of Type 1 and Type 2 prediction for geographical origin inference with SLiM simulated data. The Type 1 prediction set consists of individuals from sub-populations that are not involved in model training, whereas the Type 2 prediction set contains individuals from sub-populations partially used as a training set. Blue dots and a blue block represent individuals sampled as a training set for *Locator*. Open dots and blocks represent individuals used as prediction sets.

### Performance of *GEAplus*

To assess the performance of GEA, we conducted genome scans using simulated data filtered with a minor allele frequency (MAF) threshold of >0.01. To manage computational demands effectively, we implemented individual-based GEA with *REGENIE* and two population-based GEA approaches, as outlined below.

For individual-based GEA, we applied the same *REGENIE* settings used for the IPK landrace collection. In particular, SNPs with *r*^2^ *<* 0.2 were used for genome-wide regression to account for kinship effects, while the first three principal components of genotypic values were included as covariates to control for population structure.

For population-based GEA, sub-populations were defined based on geographical origins, followed by the calculation of allele frequencies. The sub-populations were determined by grouping individuals into geographical clusters using complete-linkage hierarchical clustering with the R function *hclust* based on geographical distances. In cases where individuals were part of the prediction set, their predicted geographical origins were used. The clustering process began by grouping individuals with pairwise geographical distances of less than 10 km. To minimize environmental heterogeneity, further subdivision of geographical groups was performed through hierarchical clustering whenever the standard deviation of the environmental variable (*SD*_env_) within a group exceeded 1% of the total *SD*_env_ across all individuals. After defining sub-populations, allele frequencies and mean environmental variables were calculated to perform the population-based GEA.

We employed two population-based GEA approaches: redundancy analysis (RDA) and a likelihood ratio test based on linear models (LM). RDA has been demonstrated to be a robust method for identifying adaptive loci with high statistical power (Forester et al. 2018). Two types of RDA were performed: simple RDA and partial RDA, the latter being conditioned on covariates. In both approaches, environmental data were used as explanatory variables, while allele frequencies served as the response variables. For partial RDA, the first three principal components of the genotypic data matrix were included as covariates to account for the confounding effects of population structure. To assess the statistical significance of SNPs, we applied chi-squared tests with 1 degree of freedom, following the framework proposed by Capblancq et al. (2018), to compute p-values.

We used four linear models for likelihood ratio tests, referred to as *LM_naive*, *LM_RR*, *LM_P*, and *LM_PRR*. The tests were conducted using our R function *geascan_rrlm*, which is available at https://gitlab.com/kjschmidlab/geaplus/-/blob/main/SLiM_gea/R_script/GEAscan_RRLMv2. R. The *LM_naive* model considered only the direct association between allele frequencies of geographical groups and environmental variation. To account for kinship and population structure, we adopted an approach similar to the individual-based GEA performed with *REGENIE*. In the *LM_P* and *LM_PRR* models, we accounted for population structure by excluding the components of environmental variation explained by the first three principal components of the genotypic data matrix. For the *LM_RR* and *LM_PRR* models, we further corrected for kinship effects by removing components of environmental variation explained by genome-wide ridge regression (RR) before performing the likelihood ratio tests.

The four models were formulated as:

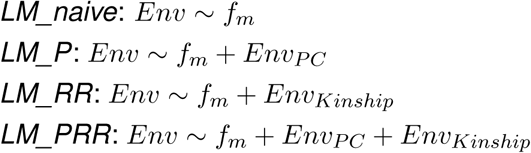

Here, *Env* represents a vector of environmental variables, and *f_m_*= {*f_m,_*_1_*, f_m,_*_2_*, f_m,_*_3_*, …, f_m,n_*} denotes the allele frequencies of *n* geographical groups at the *m*-th SNP locus. *Env_P C_* captures the environmental variation explained by the first three principal components of the allele frequency matrix *F_genotype_* = [*f*_1_*, f*_2_*, f*_3_*, …, f_m_*]. *Env_Kinship_* represents the environmental variation explained by genome-wide ridge regression.

Likelihood ratio tests were performed by calculating the test statistic:

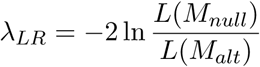

To control false positive discoveries, we adjusted *λ_LR_* using the genomic inflation factor method (François et al. 2016) and calculated p-values based on a chi-squared distribution with one degree of freedom. To compare regular GEA with *GEAplus*, we evaluated the power and true positive rate of the seven GEA approaches described above: *REGENIE*, simple RDA, partial RDA, *LM_naive*, *LM_P*, *LM_RR*, and *LM_PRR*. Regular GEA was conducted using only the training sets (*N* = 780) with true environmental variables. In contrast, *GEAplus* analyses combined the training sets (*N* = 780) with true environmental variables and the prediction sets (*N* = 14, 820), where environmental variables were derived based on predicted geographical origins.

The power of GEA was calculated as:

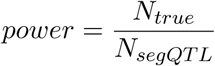

where *N_true_* is the number of detected QTLs, and *N_segQT L_* is the number of segregating QTLs. Similarly, the true positive rate (TDR) was computed as:

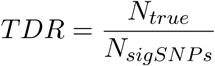

where *N_sigSNP s_* represents the total number of significant SNPs, including both neutral SNPs and QTLs.

## Results

### Prediction accuracy of geographical origins of barley landraces

We assessed the prediction accuracy (*R*^2^) of *Locator* (Battey et al. 2020) for inferring the geographical origins of IPK barley landraces using ten-fold cross-validation with ten replicates, resulting in one hundred *R*^2^ values. For data quality control, we identified 117 accessions as outliers with geo-genetic patterns that deviated from the assumption of isolation-by-distance by using *GGoutlieR*. The prediction accuracy of *Locator* was evaluated separately for the original dataset of 1,661 geo-referenced accessions and a cleaned dataset of 1,544 geo-referenced accessions using cross-validation. Through the data cleaning procedure, we improved the mean *R*^2^ values for longitude prediction from 0.932 (SD=0.039) to 0.987 (SD=0.009), and for latitude prediction from 0.839 (SD=0.062) to 0.921 (SD=0.033) (Figure 3). The range of *R*^2^ values after data cleaning also showed improvement, increasing from 0.819-0.988 to 0.954-0.996 for longitude, and from 0.637-0.947 to 0.803-0.975 for latitude (Figure 3). Furthermore, we estimated prediction errors as the geographical distances between the predicted locations and the true geographical origins. The average and median prediction errors of *Locator* models after data cleaning were 203.47 km and 84.24 km, respectively (Figure S2).

**Figure 3.**
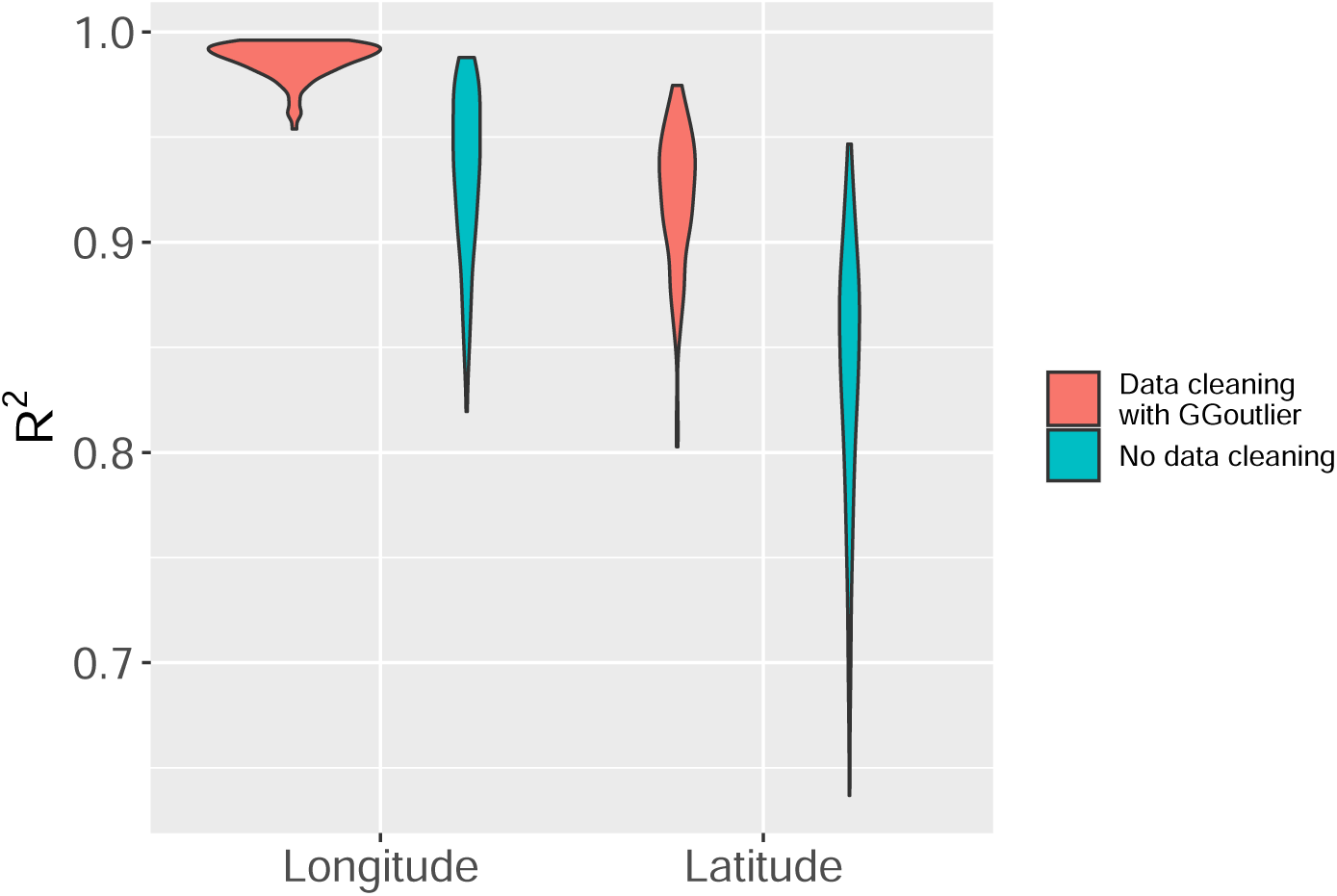
Prediction accuracy of geographical origin inference for IPK accessions. The accuracy was estimated by ten-fold cross-validation with ten replicates. The size of the population is 1,661 and 1,544, respectively, before and after data cleaning.

### GEA of barley landraces

We used the first three principal components (PCs) of 19 bioclimatic variables for subsequent GEA analyses, referring to them as environmental PC1, PC2, and PC3. Among the geo-referenced landraces, 35 out of 1,661 accessions lacked bioclimatic data because their geographical coordinates fell in regions without available data. This resulted in 1,626 geo-referenced accessions being included in the environmental PCA. Similarly, after inferring missing geographical origins, we obtained bioclimatic data for 11,032 accessions from the entire landrace collection (*N* = 12, 129) for environmental PCA. Accessions with predicted locations in regions lacking climatic data were excluded. These environmental PCs explained 79.08% and 81.69% of the environmental variance in the geo-referenced landraces (*N* = 1, 626) and the entire landrace collection (*N* = 11, 032), respectively (Figure 4 A). This indicates that the first three PCs captured the majority of the environmental variation. The PCA loadings revealed that PC1 of the geo-referenced landraces was more strongly associated with PC2 (*r* = 0.87; Table 1) compared to PC1 of the entire landrace set (*r* = 0.00; Table 1; Figure 4B and C). Using environmental PCs as response variables in *REGENIE*, we performed both regular GEA and *GEAplus*, using eleven flowering time genes as benchmarks. A flowering time gene was considered successfully identified if a significant SNP (FDR *<* 0.05) was located within a 500 kb region adjacent to the gene, based on the decay of linkage disequilibrium in domesticated barley reported in Milner et al. (2019).

**Figure 4.**
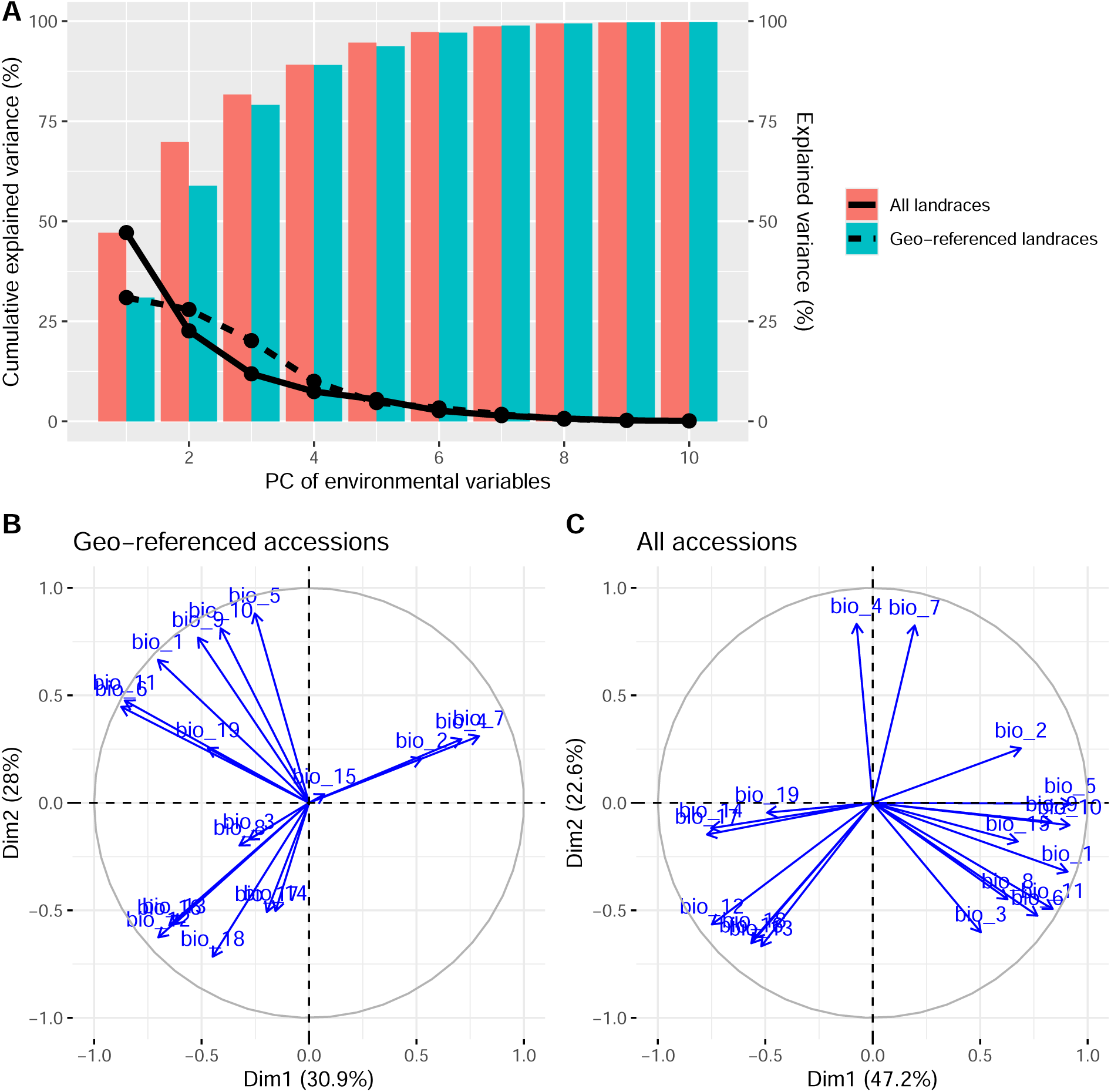
Summary of principal components (PCs) of 19 bioclimatic variables. (A) Variance explained by the first ten principal components. Colored bars represent cumulative explained variance, while lines indicate the variance explained by individual PCs. (B) Loading plot of the geo-referenced accessions (N=1,626). (C) Loading plot of the entire landrace collection with imputed environmental data (N=11,032).

**Table 1.** Correlations between the loading of the first three principal components of bioclimatic variables.

| Geo-referenced<br>landraces | All landraces |  |  |
| --- | --- | --- | --- |
|  | PC1 | PC2 | PC3 |
| PC1 | 0.00 | 0.87 | -0.37 |
| PC2 | 0.82 | 0.42 | 0.31 |
| PC3 | -0.39 | 0.18 | 0.88 |

The regular GEA analysis of IPK accessions (*N* = 1, 626) identified *PHOTOPERIOD-H1* (*PPD-H1*) on chromosome 2H and *VERNALIZATION-H1* (*VRN-H1*) on chromosome 5H as being associated with environmental PC1 (Figure 5). The most significant SNPs were located 55.6 kb downstream of *PPD-H1* and 244.3 kb upstream of *VRN-H1*, respectively. No flowering time genes were identified in the regular GEA analysis using environmental PC2 or PC3 (Figure S3). In contrast, *GEAplus* (*N* = 11, 032) identified only *PPD-H1*, and this association was with environmental PC2 (Figure 6). No flowering time genes were detected by *GEAplus* with environmental PC1 or PC3 (Figure S4). The nearest significant SNP to *PPD-H1* identified by *GEAplus* was located 225.7 kb downstream of the gene.

**Figure 5.**
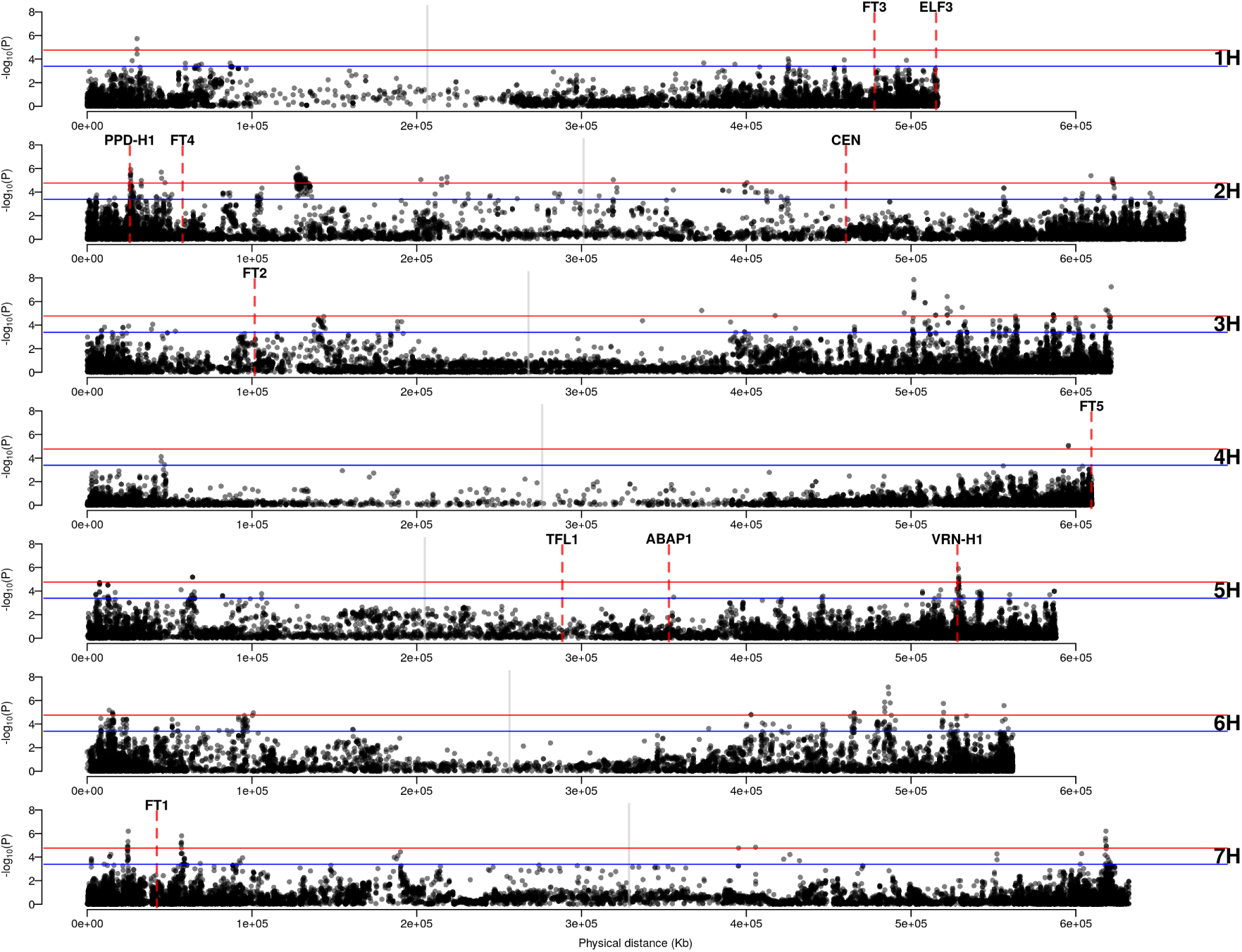
Regular GEA of IPK accessions (N=1,626) with the first environmental principal component (PC1). Blue and red horizontal lines are the significant levels of FDR = 0.05 and FDR = 0.01. Grey vertical lines indicate the positions of centromeres. Red dashed lines indicate the positions of flowering time genes.

**Figure 6.**
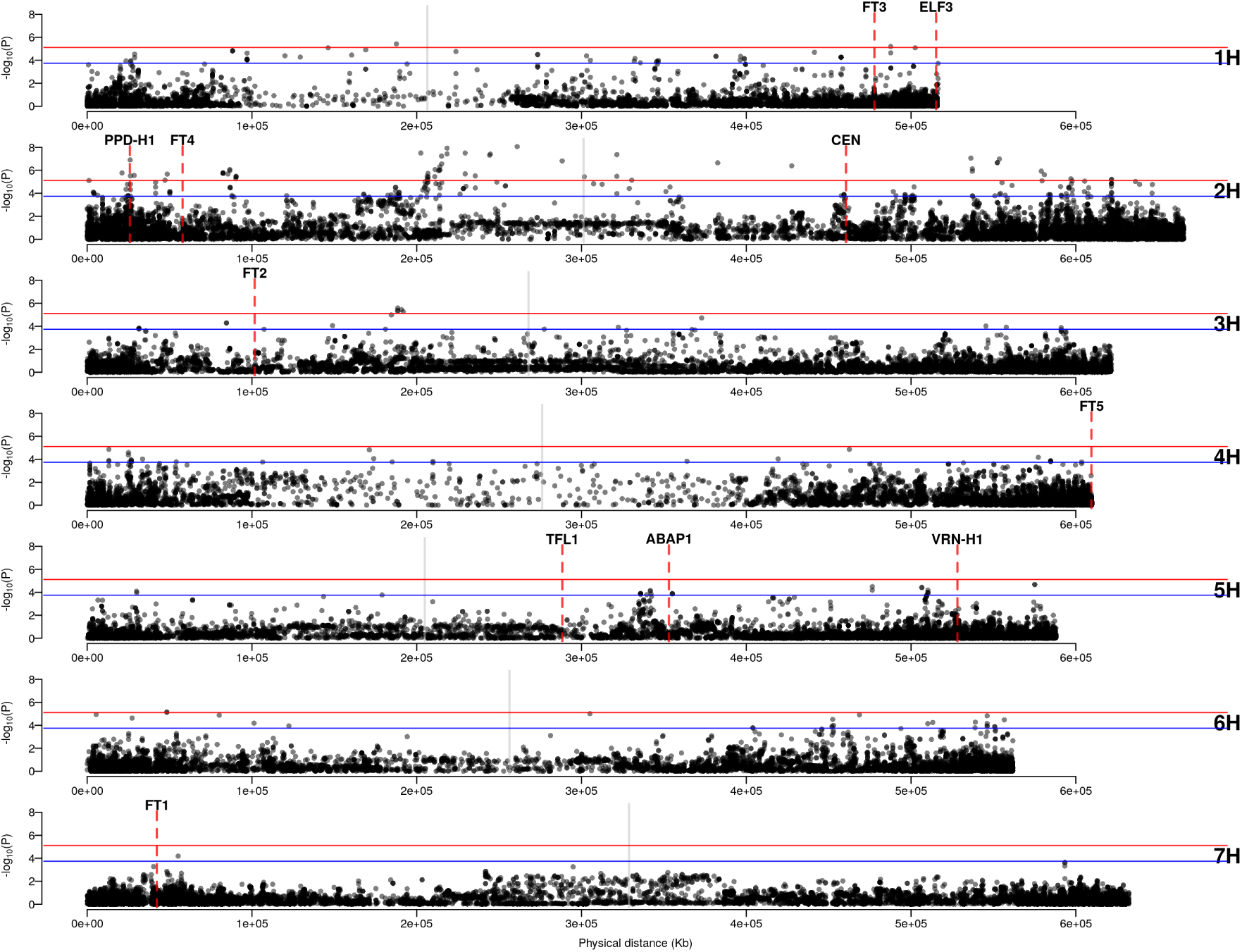
*GEAplus* of IPK accessions (N=11,032) with the second environmental principal component (PC2). Blue and red horizontal lines are the significant levels of FDR = 0.05 and FDR = 0.01. Grey vertical lines indicate the positions of centromeres. Red dashed lines indicate the positions of flowering time genes.

Comparing the two methods, regular GEA identified more significant SNPs that were closer to *PPD-H1* than those detected by *GEAplus*. Additionally, regular GEA successfully identified *VRN-H1*, which *GEAplus* did not detect. However, *GEAplus* identified four significant SNPs located 1.17 Mb upstream of *CENTRORADIALIS* (*CEN*; Figure 6), whereas regular GEA did not detect any significant SNPs near *CEN* (Figure 5, Figure S3).

### Prediction accuracy of geographical origins with simulated data

Accessions with unknown geographical origins may belong to the same sub-populations as those used to train prediction models, potentially inflating prediction accuracy due to data leakage (Bernett et al. 2024). To address this issue, we evaluated prediction accuracy using two validation approaches. Type 1 predictions introduced minimal data leakage, as the prediction set was derived from sub-populations distinct from those in the training set. In contrast, Type 2 predictions accounted for significant data leakage, with the training and prediction sets overlapping in subpopulations (Figure 2). This design reflects the realistic scenario where accessions with missing origins may not always come from sub-populations present in the training set. As expected, Type 1 predictions resulted in lower accuracy for both latitude and longitude coordinates compared to Type 2 predictions (Table 2 and Figure 7). Furthermore, Type 1 predictions exhibited standard deviations over ten times larger than those observed for Type 2 predictions (Table 2 and Figure 7). Since we used imputed environmental data in GEA, we evaluated the accuracy of predicting environmental variables based on inferred geographical origins. The imputed environmental variables exhibited strong correlations with the true environmental variables, with average *R*^2^ values ranging from 0.88 to 0.975 across four scenarios (Type 1 and Type 2 predictions paired with 1R and 2R demographic models; Figure 8). Environmental data imputation was more accurate in

**Figure 7.**
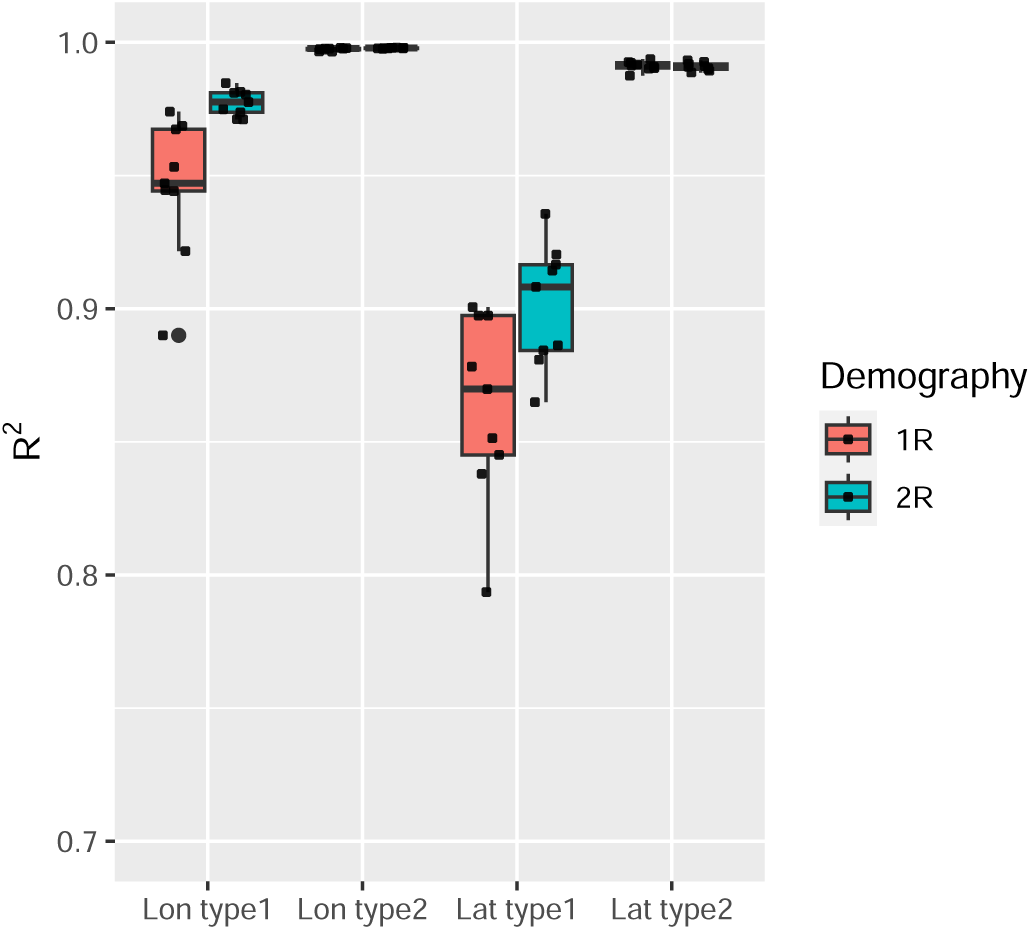
Prediction accuracy of *Locator* in SLiM simulated data. *R*^2^ is calculated between true locations and predicted locations of prediction sets separately for longitude and latitude. Prediction accuracy is evaluated with Type 1 and Type 2 prediction sets as illustrated in Figure 2.

**Table 2.** Prediction accuracy of geographical origin inference in SLiM simulated data.

| Demography | $R^2$ | | | |
| --- | --- | --- | --- | --- |
|  | Type 1 |  | Type 2 |  |
|  | Longitude | Latitude | Longitude | Latitude |
| 1R | 0.946 (0.026) | 0.864 (0.035) | 0.997 (0.0005) | 0.991 (0.002) |
| 2R | 0.977 (0.005) | 0.901 (0.023) | 0.998 (0.0001) | 0.991 (0.002) |

**Figure 8.**
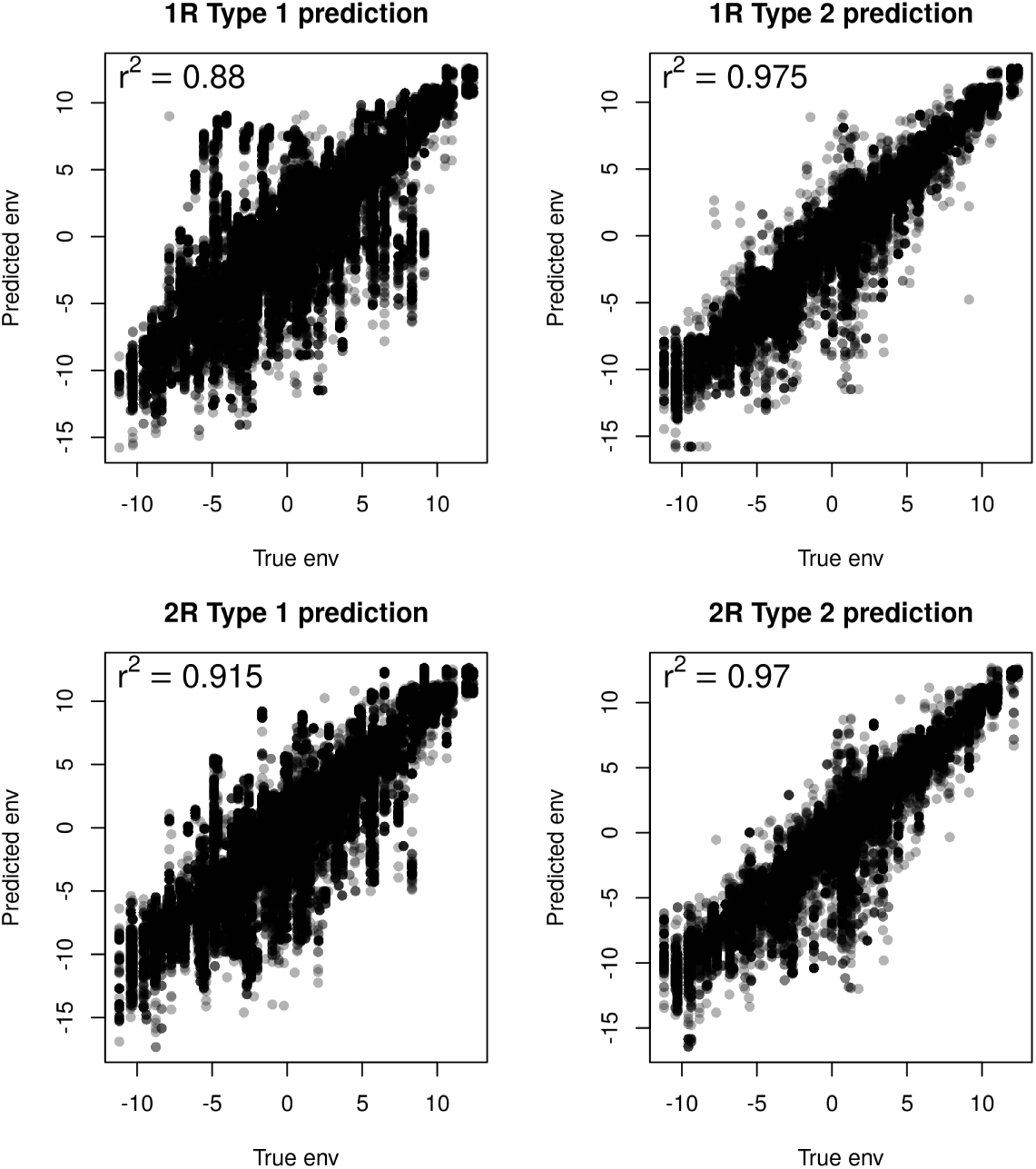
Imputation accuracy of environmental data based on predicted geographical origins in SLiM simulated data.

Type 2 predictions than in Type 1 predictions, regardless of the demographic scenario (Figure 8). Additionally, simulations of different demographic scenarios revealed that predictions were more accurate when population expansion originated from two initial sites (2R scenario) compared to a single initial site (1R scenario) (Figure 7 and Figure 8).

### Limited benefit of imputing environmental data for GEA

We tested the hypothesis that imputing environmental variables could enhance the performance of GEA by evaluating the power and true discovery rate (TDR) of regular GEA and *GEAplus*. In the simulations, QTL detection was defined as the correct identification of selected SNPs, while the detection of neutral SNPs was considered a false positive. Our results indicated that imputing environmental variables through geographical origin inference did not improve the power of GEA, regardless of the demographic scenarios or GEA approaches used (Figure 9). Both regular GEA and *GEAplus* exhibited similar performance across all tested models, except for *REGENIE*. Using population-based approaches, *GEAplus* demonstrated power comparable to regular GEA. However, with the individual-based GEA approach using *REGENIE*, regular GEA exhibited higher power than *GEAplus* in both the 1R and 2R demographic scenarios (Figure 9A and B). This finding contradicts our hypothesis that imputing environmental data would enhance the identification of adaptive loci with GEA.

**Figure 9.**
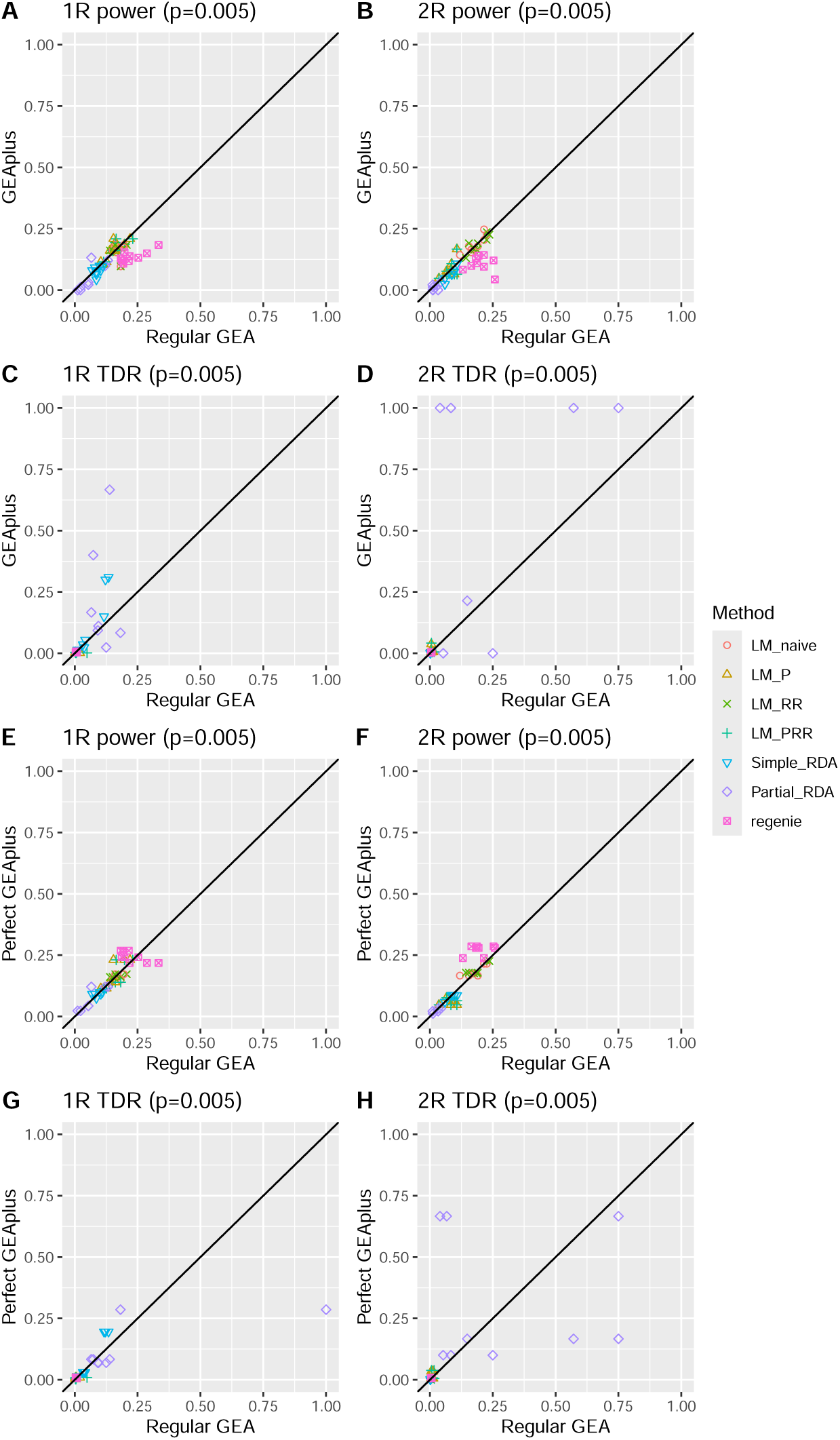
Comparison of power and true discovery rate (TDR) between regular GEA (*N* = 780) and *GEAplus*/*perfect GEAplus* (*N* = 15, 600). (A-D) Pairwise comparisons between regular GEA and *GEAplus* under 1R and 2R demographic scenarios. Power and TDR are defined as the proportion of detected QTLs (*p* = 0.005) relative to the total number of segregating QTLs and the number of significant SNPs, respectively. TDR of *GEAplus* with RDA is missing in four simulations, two with 1R setting and two with 2R setting, because of no significant SNP. (E-H) Pairwise comparisons between regular GEA and *perfect GEAplus* under 1R and 2R demographic scenarios. *Perfect GEAplus* assumes environmental information is imputed with no error.

We also evaluated the true positive rate (TDR) to assess the accuracy of correctly identifying adaptive loci. Simple RDA and partial RDA showed a tendency toward higher TDR in *GEAplus* compared to regular GEA, while other approaches did not exhibit this pattern (Figure 9C and D). However, *GEAplus* with partial RDA failed to identify any significant SNPs in two of the simulation runs for both 1R and 2R scenarios, resulting in missing TDR values, which are not shown in Figure 9C and D.

In addition to investigating *GEAplus*, we examined a hypothetical scenario of perfect imputation of environmental variables, referred to as *perfect GEAplus*. To simulate this condition, we used the true environmental variables to conduct GEA on the entire simulated population and evaluated the power and TDR of *perfect GEAplus*. We found that the power of *perfect GEAplus* was not significantly different from that of regular GEA in the 1R scenario across all GEA approaches (Figure 9E). However, in the 2R scenario, individual-based GEA using *REGENIE* demonstrated slightly higher power compared to regular GEA (Figure 9F). Among all approaches, individual-based GEA with *REGENIE* exhibited the highest power in both the 1R and 2R scenarios (Figure 9E and F). When comparing TDR, no significant differences were observed between *perfect GEAplus* and regular GEA (Figure 9 G, and H). However, population-based partial RDA tended to achieve a higher TDR compared to other approaches in our simulations, particularly in the 2R scenario (Figure 9G and H). Overall, our findings suggest that increasing the sample size does not significantly enhance the performance of GEA under our simulated scenarios, consistent with our empirical analysis.

## Discussion

Our approach of integrating imputation of environmental data and GEA rests on two key assumptions: first, that genomic sequences can accurately predict missing geographical origins based on the isolation by distance expectation, i.e. the majority of genetic variation are neutral and reflect demographic history instead of adaptation (Wright 1943), and second, that GEA can achieve higher power with increased sample sizes (Lotterhos and Whitlock 2015). While we observed good accuracy in geographical origin inference, the significant expansion in sample size did not translate into substantial benefits for GEA.

### Performance of geographical origin inference

Using genome-wide sequencing data, we employed the deep-learning-based geographical origin inference approach *Locator* (Battey et al. 2020) to recover missing location data for over ten thousand accessions. Our cross-validation results showed high accuracy (*R*^2^ *>* 0.9) in predicting both longitude and latitude. We also evaluated the accuracy of geographical origin prediction using forward genetic simulations with a scaling approach to ensure computational feasibility. A recent study highlighted the need to increase the number of burn-in iterations in forward simulations, depending on the effective population size and sequence length, to obtain unbiased genetic diversity when using the scaling approach under the Wright-Fisher model (Ferrari et al. 2024). However, our study used the *nonWF* model of SLiM (Haller and Messer 2019) and focused on GEA rather than genetic-diversity-based metrics, making reduced genetic diversity due to scaling less of a concern. Our simulations revealed that *Locator* performed better under a population expansion scenario originating from two locations (2R scenario) compared to a single origin (1R scenario; Table 2). This suggests that the presence of population genetic structure may enhance the accuracy of geographical origin inference using neural networks. Additionally, we observed that prediction accuracy was higher for individuals derived from sub-populations overlapping with the training set (Type 2 prediction set) than for those sampled from entirely excluded sub-populations (Type 1 prediction set; Table 2). This finding indicates that prediction accuracy can vary depending on the genetic similarity between tested samples and the individuals in the training set. To optimize the training set, the ideal sampling strategy should include as many sub-populations as possible, even if the sample size from each sub-population is small.

In our empirical analysis of genebank accessions, cross-validation revealed that the accuracy of *Locator* was strongly influenced by the extent to which the training set adhered to the isolation-by-distance assumption. We observed a significant improvement in prediction accuracy after removing outliers that violated this assumption. This finding highlights the critical role of data quality and population history in accurately inferring geographical origins. Populations with a history of recent long-distance migration events pose a particular challenge for geographical origin prediction. For instance, if migrants recently established a large colony in a distant environment, the deep-learning approach *Locator* may incorrectly infer geographical origins due to high genetic similarity between the original and newly established populations. Such scenarios are plausible for plant species that have dispersed through human activities or seed exchange in recent decades as postulated for Ethiopian barley traditional cultivars (Teklemariam et al. 2022). Therefore, recovering missing geographical origins from genotypic data requires a solid understanding of population history and careful data preprocessing to ensure reliable predictions.

#### Improving geographical origin inference with ecological knowledge

In our cross-validation of geographical origin inference for barley landraces, we observed some predictions that were ecologically implausible. The deep learning approach, *Locator*, minimizes prediction error by reducing the distance between predicted and true locations. However, it does not account for ecological constraints, such as whether the predicted locations are suitable for barley growth. For example, in our analysis of the barley landrace collection, 1,097 accessions were erroneously projected to the Mediterranean Sea, resulting in failed environmental data imputation. These errors stem from the limitations of the current deep learning model, which attempted to predict geographical origins for samples that might represent an admixture of landraces from Europe and North Africa. Additionally, the model overlooks the impact of microclimates, such as those caused by altitude in mountainous regions, where climate conditions can vary significantly over short distances and lead to distinct evolutionary outcomes (Hämälä and Savolainen 2019; Lampei et al. 2019). These limitations present a significant obstacle to using geographical origin inference in GEA studies. Inaccurate environmental data imputation, even when geographical origin inference achieves high accuracy, can introduce substantial noise into GEA analyses.

To improve the quality of environmental data imputation, incorporating knowledge of natural habitats is essential. A recent study proposed a statistical framework to estimate the credibility of individual geographical records for hundreds of species (Arlé et al. 2021). This framework assesses the likelihood of geographical origin records being accurate by leveraging species distribution data from biological databases. Integrating such probability-based plausibility measures into the training process of deep learning models could significantly enhance the reliability of geographical origin inference.

### Effect of imputing environmental data on GEA

Using imputed environmental data, GEA has the potential to uncover distinct adaptive signatures, although this does not necessarily translate into increased power for genome-wide scans. In our implementation of *GEAplus* on barley landraces, we defined signals near flowering time genes as successful identifications of adaptive loci. Both regular GEA and *GEAplus* identified significant signals near *PPD-H1*, a gene associated with adaptation to photoperiod, temperature, and drought (Gol et al. 2021; Ochagavía et al. 2022; Turner et al. 2005; Wiegmann et al. 2019). Recent research involving the re-sequencing of *PPD-H1* in a global germplasm collection revealed associations between *PPD-H1* haplotypes and precipitation-related bioclimatic variables (Sharma et al. 2020). In contrast to the consistent signatures near *PPD-H1*, no significant SNPs close to *VRN-H1* were found with *GEAplus*, while new SNPs emerged upstream of *CEN*(Figures 5 and 6). Although the possibility of false-positive discoveries cannot be ruled out, this observation may reflect the impact of geographical diffusion of mutations on GEA approaches. The efficacy of GEA depends on the differentiation of allele frequencies across distinct environments. Non-monotonic clinal patterns in allele frequencies along environmental gradients can result in the failure to detect adaptive loci using GEA (Lotterhos 2023). Moreover, the spread of mutations is often constrained by geographical distance, limiting adaptive alleles to specific areas. This can lead to non-clinal patterns where adaptive alleles are absent in geographically distant regions with comparable environmental conditions. Such scenarios are plausible in global germplasm collections, where accessions may originate from geographically distant yet ecologically similar environments.

We can conclude that environmental data imputation can influence GEA by either revealing new adaptive loci or eliminating previously observed adaptive signatures due to changes in allele frequencies. This issue is not unique to the *GEAplus* framework but also applies to regular GEA when samples are collected across a broad geographical range, as monotonic clinal patterns are a prerequisite for the success of GEA (Lotterhos 2023). To address this limitation, one potential solution is to perform multiple GEA tests within pre-defined smaller geographical areas using a sliding window approach, rather than conducting a single GEA test with all samples combined. This localized approach may improve the identification of adaptive loci by uncovering region-specific variations. However, this method introduces a large number of statistical tests, as it requires evaluating tens of thousands of SNPs for each environmental variable across hundreds of geographical windows. Such extensive testing could lead to a proliferation of spurious signals. Therefore, further development and rigorous validation of this methodology are crucial to ensure its effectiveness and reliability.

### Validation of GEA approaches by simulations

In our comparison between *GEAplus* and regular GEA using simulations, we observed a slight improvement in power for individual-based GEA with *REGENIE* through environmental data imputation in the 2R scenario, but no such improvement for population-based GEA approaches, even with perfectly imputed environmental data (Figure 9F). This finding is consistent with observations reported by Lotterhos and Whitlock (2015), who noted that increasing the sample size improved the power of LFMM (Frichot et al. 2013), an individual-based method, but not Bayenv2.0 (Günther and Coop 2013), which is population-based. The difference in power was attributed to the individual-based versus population-based units of analysis in these approaches. Despite differences in simulation scenarios, our results align with Lotterhos and Whitlock (2015), suggesting that individual-based GEA tends to achieve higher statistical power than population-based GEA when sample sizes are large. Aside from the slight power improvement observed for the individual-based approach in the *perfect GEAplus* of the 2R scenario, our simulations indicated no general advantage of *GEAplus* over regular GEA, even with perfectly imputed environmental data (Figure 9E, F, G, and H). Previous simulation studies have suggested that GEA approaches are generally insensitive to sample size, except under island model scenarios (Forester et al. 2018; Lotterhos and Whitlock 2015). Since our simulations approximated an isolation-by-distance demographic structure in a confined geographical region, increasing the sample size did not significantly benefit GEA performance, as also observed in earlier studies (Forester et al. 2018; Lotterhos and Whitlock 2015). This was true even though the sample size in the *GEAplus* scheme was twenty times larger than that in the regular GEA scheme. However, if GEA were applied on a continent-wide scale with sufficient isolation between populations, larger sample sizes could potentially improve performance. In addition to demographic effects, various other factors, such as mating systems (Hodgins and Yeaman 2019), genome recombination rates (Lotterhos 2019), and sampling strategies (Lotterhos and Whitlock 2015), can influence the detection of adaptive loci. Future research incorporating diverse simulation settings that account for these factors will be essential to gain a more comprehensive understanding of the conditions under which GEA performs effectively.

## Conclusion and outlook

In summary, our study demonstrates the potential of using extensive genomic data and deep learning models to recover missing geographical origins in global germplasm collections, providing ecological insights for identifying alleles associated with tolerance to environmental stresses. However, we also acknowledge the limitations of current geographical origin inference methods. Existing neural networks prioritize minimizing geographical distance prediction errors without considering the ecological plausibility of the predicted habitats. Our findings suggest that environmental data imputation primarily benefits individual-based GEA. Additionally, our simulations indicate that large-scale genotyping is unlikely to improve GEA performance if samples are indiscriminately pooled for analysis. Instead, increasing the volume of data is more likely to enhance the detection of region-specific associations rather than global GEA patterns. The development of innovative approaches, such as an individual-based sliding window GEA method, will be crucial for effectively leveraging imputed environmental data while accounting for the geographical distribution of alleles.

## Acknowledgments

We sincerely thank Dr. Martin Mascher and Max Haupt of the Leibniz Institute of Plant Genetics and Crop Plant Research (IPK) for providing the raw VCF data of barley landraces used in this study. This research was supported by funding from the Federal Ministry of Food and Agriculture (BMEL) through the Federal Office for Agriculture and Food (BLE) under the Federal Programme for Ecological Farming and Other Forms of Sustainable Agriculture (Project number 2818202615). Additional support for C.W.C. was provided by the Study Abroad Fellowship from the Ministry of Education, Taiwan (R.O.C.) (Project number 1100123625).

## Data archiving

The scripts used for data analysis and simulations in this study are available at https://gitlab. com/kjschmidlab/geaplus. The VCF file of IPK barley landraces will be made available upon request.

