## Supplementary Text and Figures for "Effects of using deep learning to predict the geographic origin of barley genebank accessions on genome-environment association studies"

### SLiM simulation

#### Mutational effect of QTLs

In our simulation, we assumed that the environmental variables of all sites are located in a 95% interval of the expected genetic variation. The expected genetic variation is calculated as $\sigma_g^2 = 2N\sigma_{qtl}^2$, where $N$ is the number of QTLs ($N = 100$) and 2 is for diploidy. Let the boundary of the 95% interval of the expected genetic variation be $\pm c\sigma_g$, where $c$ is a constant, and let the most extreme environmental variable after centering be $x$. With our assumption, $|x|$ should be equal to or less than $c\sigma_g$. Therefore, we have $c\sigma_g \geq |x|$. We can rewrite it as $c\sqrt{2N\sigma_{qtl}^2} \geq |x|$, and it gives $\sigma_{qtl} \geq \sqrt{\frac{|x|^2}{2Nc^2}}$. With the equation above, we set $\sigma_{qtl} = 0.45$.

#### Plasticity of selection

We determined the plasticity, standard deviation (SD) of fitness bell curve, based on the environmental contrasts between connected sites. This was done by calculated the absolute difference between the environmental variables of the two connected sites, denoted as $C_{env} = |Env_i - Env_j|$, where site $i$ is a connected neighbor of the site $j$ with gene flow.

To simulate a sufficiently strong isolation by environment, we assumed that 90% of $C_{env}$ values fall within a 95% interval of the fitness bell curve. Among the selected 312 sites, we found that 90% of $C_{env}$ values were less than 5.7. To set the plasticity parameter ($\sigma_{plasticity}$) for the SLiM simulation, we chose a value of 2.85, such that $2\sigma_{plasticity} = 5.7$, approximately covering the 95% interval under a normal distribution.

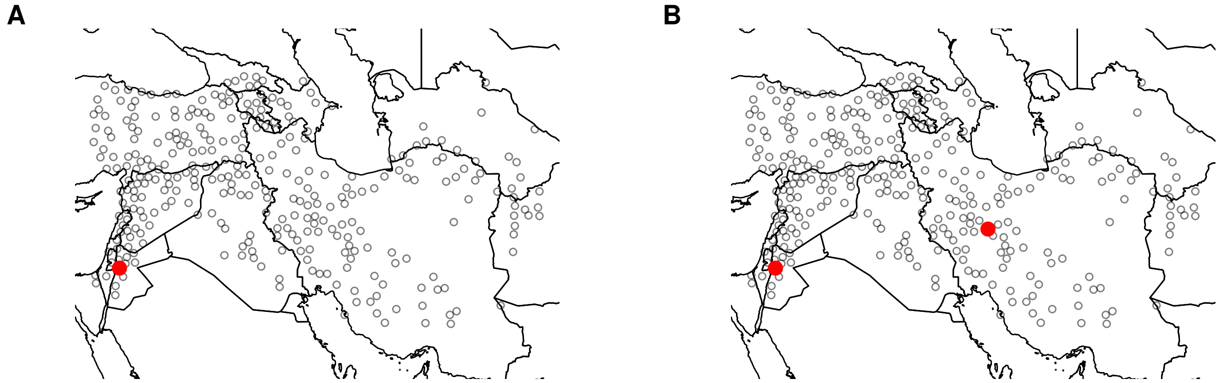


**Figure S1** Geographical distribution of sub-populations in two demographic scenarios of SLiM simulation. A. Population expansion from one refugium (1R). B. Population expansion from two refugium (2R). Red dots indicate the starting points of population expansion.

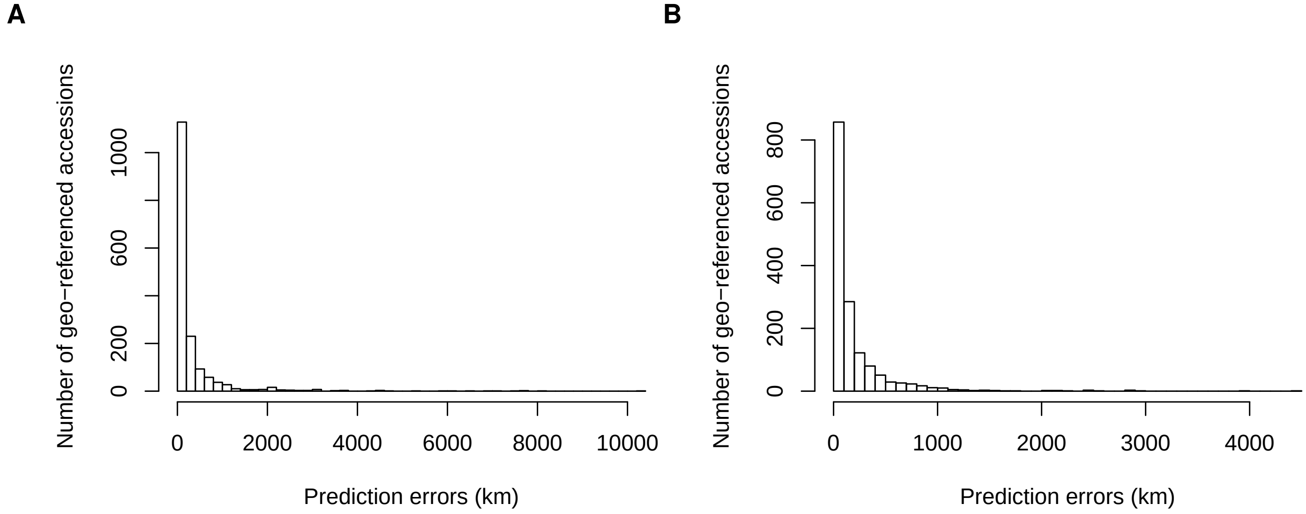


**Figure S2** Mean prediction errors in kilo-meters estimated from cross-validation of IPK barley landrace collection. A. Prediction errors of original samples. B. Prediction errors of samples excluding outliers with unusual geo-genetic patterns.

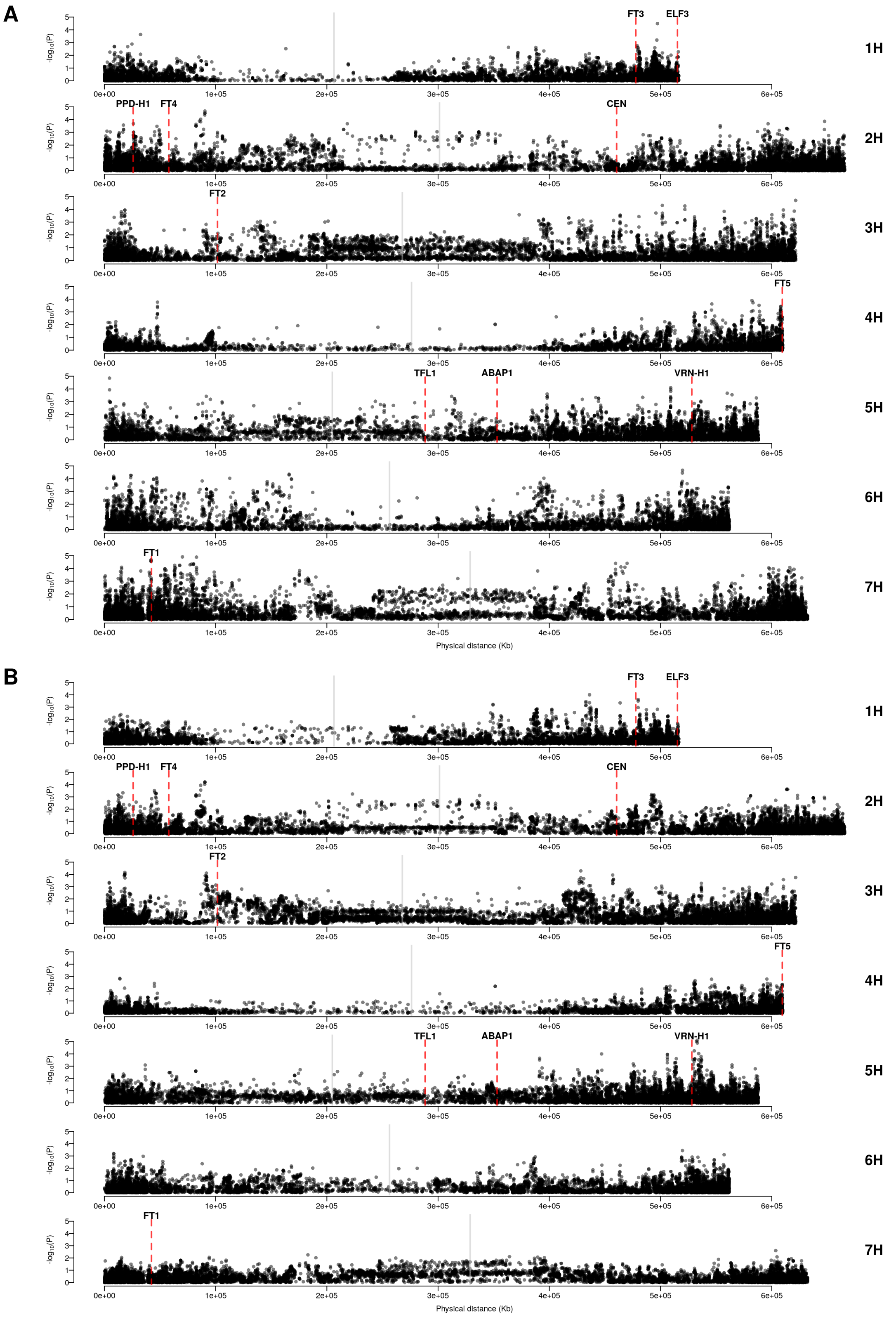


**Figure S3** Regular GEA of IPK landraces with the environmental principal component (PC). A. Regular GEA with environmental PC2. B. Regular GEA with environmental PC3. Blue and red horizontal lines are the significant levels of FDR = 0.05 and FDR = 0.01. Grey vertical lines indicate the positions of centromeres. Red dashed lines indicate the positions of flowering time genes.

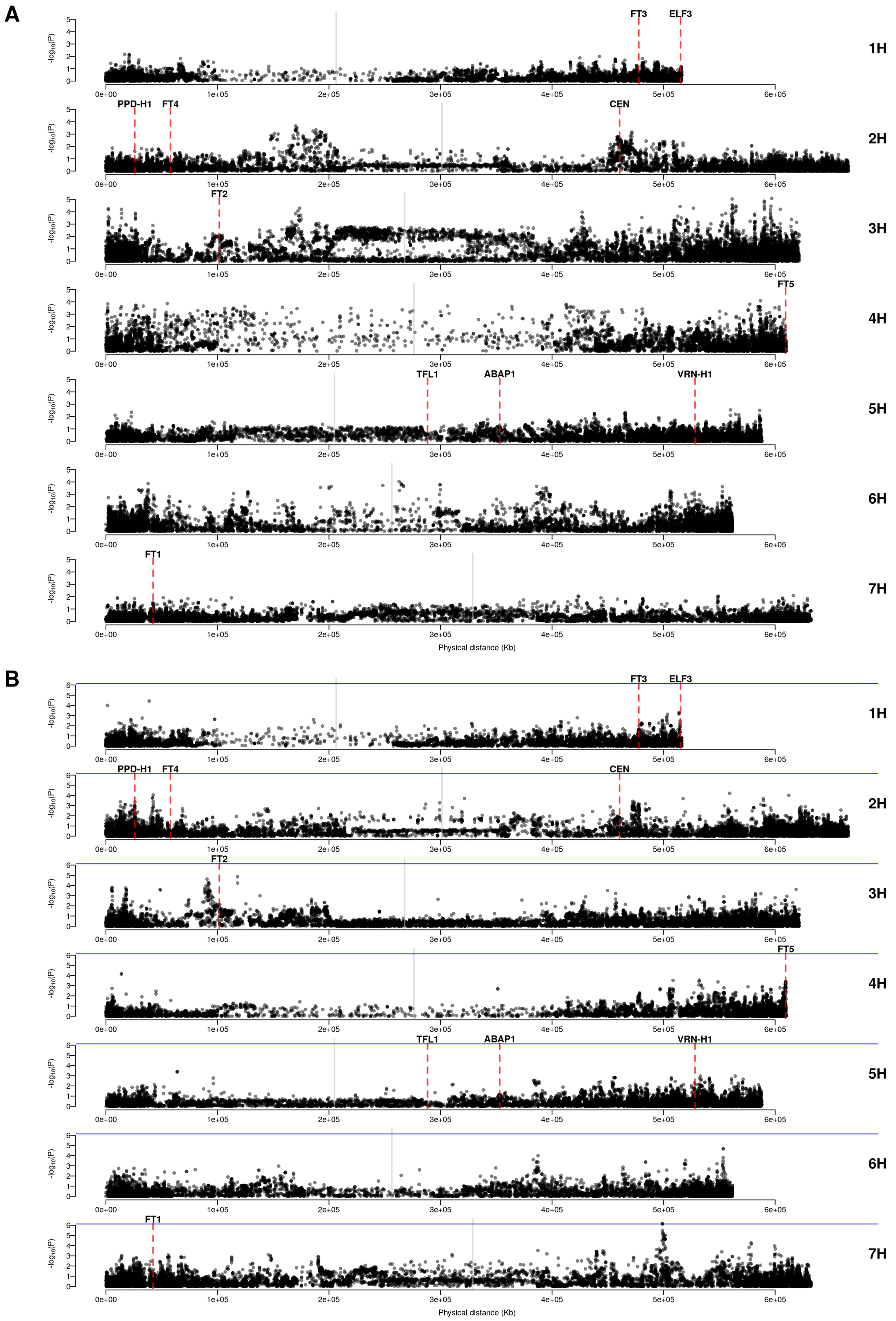


**Figure S4** *GEAplus* of IPK landraces with the environmental principal component (PC). A. *GEAplus* with environmental PC1. B. *GEAplus* with environmental PC3. Blue and red horizontal lines are the significant levels of FDR = 0.05 and FDR = 0.01. Grey vertical lines indicate the positions of centromeres. Red dashed lines indicate the positions of flowering time genes.
